# Transcription driven repurposing of cardiotonic steroids for Lithium treatment of severe depression

**DOI:** 10.1101/2025.01.24.634703

**Authors:** Richard Killick, Claudie Hooper, Cathy Fernandes, Christina Elliott, Dag Aarsland, Svein R. Kjosavik, Ragnhild Djønne Østerhus, Gareth Williams

## Abstract

Lithium is prescribed as a mood stabiliser in bipolar disorder and severe depression. However, the mechanism of action of Lithium is unknown and there are major side effects associated with prolonged medication. This motivates a search for safer alternative drug repurposing candidates.

Given that the drug mechanism may be encoded in transcriptional changes we have generated the gene expression profile for acute Lithium treatment of cortical neuronal cultures. We found that the Lithium associated transcription response harbours a significant component that is the reverse of that seen in human brain samples from patients with Major Depression, Bipolar Disorder and a mouse model of depression. Interrogating publicly available drug driven expression data we found that cardiotonic steroids drive gene expression in a correlated manner to our acute Lithium profile. An analysis of the psychiatric medication cohort of the Norwegian Prescription Database showed that cardiotonic prescription is associated with a lower incidence of lithium prescription.

Our transcriptional and epidemiological observations point towards cardiotonic steroids as possible repurposing candidates for Lithium. These observations motivate a controlled trial to establish a causal connection and genuine therapeutic benefit in the context of depression.

## Background

Despite the long therapeutic pedigree of Lithium (Li) as a mood stabiliser[1] its mechanism of action is still unclear and use is associated with major side-effects[2]. The side-effects are likely due to the non-specific effects of Li covering the modulation of serotonin[3], dopamine[4], and glutamate[5] neurotransmitters the inhibition of glycogen synthase kinase-3 (GSK-3)[6] and inositol monophosphatase[7, 8]. Li can also enter through sodium channels[9], block potassium channels[10] and impact calcium mobilisation[11] and subsequently modulates neuronal ion channel activity and membrane potential[12]. The side-effect burden[2] motivates a search for an alternative medication and the wide spectrum of activity points towards a candidate emerging based on a global phenotype comparison. In particular, a drug repurposing strategy, where existing therapeutics emerge as candidates for conditions for which they weren’t originally developed, is a potential avenue based on the availability of extensive activity data from laboratory experiments and prescription histories[13]. One insight into the mechanism can be obtained via global transcription profiling[14-16]. A direct validation of drug associated transcription encoding therapeutic activity would be the observation that the gene expression changes driven by the drug are reverse of those seen in disease states[17, 18]. This, transcription-based drug repurposing approach, has been used in the search for alternative candidates in cancer therapy[17, 19] and neurodegenerative disease[13, 20, 21]. Mapping biological activity to transcriptional perturbation is a particularly fertile option as the data is both quantitative and multidimensional with an abundance of publicly available data on disease state profiles, through online depositories such as the National Center for Biotechnology Information (NCBI) Gene Expression Omnibus (GEO)[22], and drug associated profiles via the Connectivity Map (CMAP)[14] and Library of Integrated Network-Based Cellular Signatures (LINCS)[23] datasets. One can also base repurposing on actual epidemiological data where disease incidence is tracked in relation to drug candidate use. A recent notable example is the repurposing of salbutamol for Parkinson’s disease (PD)[24], where the incidence of PD over a five year period was shown to decrease in proportion to salbutamol, a beta-adrenergic agonist, prescription and increase with the prescription frequency of a beta-adrenergic antagonist, propranolol. The obvious drawback of this methodology is that retrospective study analysis can at best point to causality and serve rather to either confirm hypotheses or motivate further investigation or controlled trials. In the present work we sought to delimit repurposing candidates for Li on the basis of an expression profile correlation and go some way to validate the candidates based on a correlation analysis of prescription data, with the hypothesis that those taking the candidate drug are less likely to be prescribed Li.

We first sought to define an activity profile for Li. We reasoned that the most likely activity representation would be furnished by gene expression profiling, as there is a wealth of data on compound driven gene expression changes[14, 23]. We also wanted to focus on the immediate downstream effects of Li treatment as these will more likely reflect the drug’s direct engagement with primary targets and avoid confounding effects due to compensatory changes. Assuming the therapeutic effects are primarily driven by neuronal interactions, we profiled Li in primary neuronal cultures. We found that our acute Li profile (ALP) shows a significant negative correlation with transcriptional profiles corresponding to major depressive disorder (MDD) and bipolar disorder (BP). Interestingly, we also found a negative correlation with a chronic variable stress (CVS) mouse model brain profile. This observation bolstered the hypothesis that ALP captures some of the therapeutic aspects of Li activity and motivated our search for compounds with similar transcriptional profiles as possible repurposing candidates. We found that multiple cardiotonic steroids (CTS) significantly correlate with the ALPS, MDD and CVS profiles.

There are two CTS currently in the clinic: digoxin and digitoxin. We reasoned that if CTS use has a therapeutic component in common with Li then there may be fewer prescriptions of Li in those on CTS medication. To investigate this possibility, we gathered data from the Norwegian Prescription Database (NorPD, www.norpd.no) that consists of prescription data across the public health sector from 2004 onwards. We decided to focus our analysis exclusively on individuals who have been prescribed at least one form of psychiatric medication, thus aiming to better isolate the specific factors influencing the decision to prescribe lithium. This restriction ensures that our study population has already engaged with psychiatric care at a level that necessitates pharmacological intervention, hence providing a more relevant and homogeneous baseline for comparison. Including individuals without any psychiatric medication history might introduce confounding variables related to barriers to care, varying diagnostic thresholds, or treatment-seeking behaviors, which could dilute or obscure the factors specifically associated with lithium prescription. This cohort still comprises a significant fraction of the entire population (2 million of 5.5 million). To this end we compiled prescription records for 94 psychiatric medications and two CTS (digoxin and digitoxin) for the entire Norwegian population covering the years 2010 to 2021. Our analysis points to a significant reduction in Li prescriptions for those on CTS medication.

## Methods

### Cell culture treatment

All materials were obtained from Sigma (UK) unless otherwise specified. Time-mated Sprague Dawley rats were from Charles River (UK). Neurobasal medium, B27 supplement and Trizol were from Invitrogen (UK). Papain dissociation systems were from Worthington (UK). Streptavidin-phycoerythrin was from Molecular probes (UK); Biotinylated anti-streptavidin antibody from Vector Laboratories (UK); Taqman reverse transcription (RT) reagent kit from Applied Biosystems (UK); QuantiTect SYBR green kit from Qiagen (UK) and Rat Genome 230 v2.0 whole genome microarray chips and the one cycle target labeling and control kit were from Affymetrix (UK).

Primary cortical neuronal cultures were generated from Sprague Dawley E18 rat embryos using papain dissociation according manufacturers’ instructions. Neurons were cultured in neurobasal medium supplemented with B27 for 7 days at 37°C in a humidified atmosphere of 5% CO_2_ in air as previously described[25]. After 7 days in culture (d.i.c.) neurons were washed twice in neurobasal medium (without any supplements) then left in fresh neurobasal medium for 2 h, this to remove insulin, a constituent of B27 supplement and an inhibitor of GSK3. Following this incubation period LiCl was then added to the media at a final concentration of 10 mM and cells cultured for a further 2 h before collection.

### Microarray data

Total RNA was extracted from primary rat cortical neurons 2 h post treatment using Trizol followed by isopropanol precipitation. The RNA integrity number (RIN), a measure of RNA quality (10 being maximal), ranged from 9.6-10 for all samples, as determined using an Agilent Bioanalyser. Total RNA (7.5 mg) was reversed transcribed using oligo (dT), biotinylated and hybridized to Affymetrix Rat Genome 230 v2.0 whole genome microarrays using the one cycle target labeling and control kit according to protocols in the Affymetrix GeneChip Expression Analysis Technical Manual. The arrays were washed and stained with streptavidin-phycoerythrin and the fluorescent signal amplified with biotinylated anti-streptavidin antibody. Staining and washing were performed using an Affymetrix fluidics system and microarrays scanned using an Affymetrix gene chip 3000 scanner. The full expression profile for Li is given in Supplementary Table S2.

### Expression profile analysis

The CEL file data was RMA normalized using the affy package in the Bioconductor R environment (http://www.bioconductor.org/). Differential gene expression profiles for the lithium treatment done in our laboratory and for the data sourced from NCBI GEO were generated using the R limma package. To facilitate expression profile comparison and generate composite profiles the gene expression changes were converted to Z score values and composite profiles defined as the summed Z scores divided by the square root of the number of experiments for each gene.

Repurposing candidates were defined based on a global transcription correlation analysis where the query profile was scored against the CMAP[14] repository of drug driven differential expression profiles, consisting of 1,309 drug-like compounds.

### Publicly available MDD and BPD gene expression data

Interrogating the NCBI GEO gene expression repository, we found five expression series corresponding to brain samples from sufferers of Major Depressive Disorder (MDD) with matched controls with GEO identifiers GSE101521[26], GSE102556[27], GSE53987[28], GSE44593 and a study[29] split over the following series: GSE54562, GSE54563, GSE54564, GSE54565, GSE54567, GSE54562, GSE54568, GSE54570, GSE54561, GSE54572, GSE54575. This enabled us to delimit an MDD meta-profile based on combining the Z scores for gene expression changes relative to control groups from the various independent data sets. In cases where donor measures were included in the data, such as age, sex, brain region, suicide status, these were factored into the differential expression analysis by being included as covariates in the linear model. Similarly, we generated a Bipolar Disorder (BPD) meta-profile from six independent series (GSE5392[30], GSE53239[31], GSE53987[28], GSE62191[32], GSE80336[33], and GSE81396[34]). Our metanalysis also included a differential expression profile for a mouse chronic unpredictable stress (CUS) model of depression[35] GSE102556[27]. See Supplementary Table S3 for the summary profiles.

### NorPD prescription data

Prescription data consisting of prescription date, date of birth, date of death in case of death were obtained from the NorPD prescription database for 95 psychiatric medications (802,109 men and 1,123,759 women) and two commonly prescribed cardiotonics, digoxin and digitoxin (24,099 men and 23,996 women and 14,994 men and 17,890 women of these in the psychiatric medication cohort), see Supplementary data Table S1 for the Anatomical Therapeutic Chemical (ATC) codes.

## Results

### Transcriptional changes associated with acute Li treatment

On the premise that knowing which genes are responsive to Li could help elucidate the mechanism of action we examined the transcriptional effects of Li on rat primary cortical neurons using whole genome expression microarrays. A number of microarray-based expression studies, performed in model systems ranging from yeast through neuronal cell lines to brains of treated rodents, have been reported. A considerable number of gene expression studies have been undertaken in varied model systems with the aim of shedding light on the mechanism of action of Li on mood[36-41]. All employed chronic treatment regimens given Li requires from several weeks to up to six months to show maximal clinical benefit in man. We reasoned that such chronic treatments could miss the acute, “first wave”, effects of Li on gene expression due to masking by subsequent waves of transcription. In the light of this we measured the transcription changes two hours post treatment of rat cortical neuronal cultures.

We found that a disproportionate number of the genes regulated by the acute treatment encode for transcription factors, thus we believe we have uncovered, for the first time, the direct effects of Li on neuronal gene transcription. In total 2157 genes (643 UP 1514 DOWN) showed expression changes at the p < 0.05 significance level, see Figure 1 for a list of the most highly regulated genes. In addition to the regulation of transcription factors notable amongst the up-regulated genes implicated in neuronal function are: the brain derived neuronal growth factor (BDNF), all three members of the nuclear receptor subfamily 4A (NR4A) of genes involved in synaptic mitochondrial function, and calcium dependent kinase kinase 2 (CAMKK2) involved in learning and memory.

**Figure 1.**
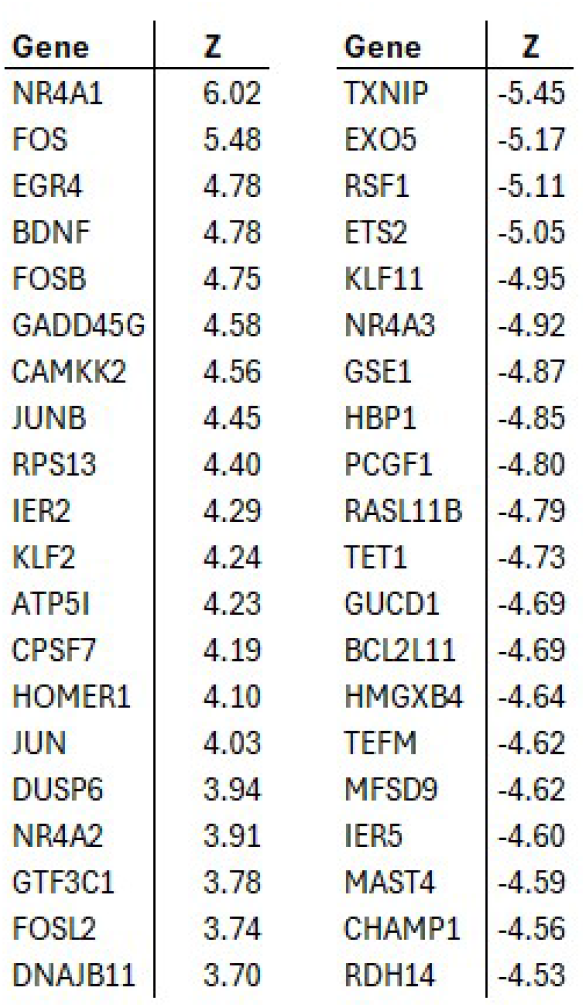
Acute Lithium expression response. Lithium elicits a substantial transcriptional response two hours post treatment of cortical neurons differentiated from embryonic rat progenitor cells. In total 2157 genes (643 UP 1514 DOWN) showed expression changes at the p < 0.05 significance level.

### The Li expression profile in the context of depression

It is known that depression is associated with consistent expression changes in the brains of sufferers, see for example Pantazatos SP *et al* [26] and this motivated us to investigate the relationship of depression associated expression changes with our acute Li expression profile. Our hypothesis being that if a component biological activity is encoded in the perturbation of gene expression we would expect the signature of a therapeutic to have a component that is reverse of that seen in the corresponding disease state. Interrogating the NCBI GEO repository, we found five patient cohorts corresponding to brain sample from sufferers of MDD, see Methods for details. It is apparent that the genes most responsive to Li treatment tend to be perturbed in the reverse sense in MDD and to a lesser extent in BPD, see Figure 2. Of interest is that a chronic variable stress (CVR) mouse model of depression shows a consistent expression profile with MDD. The mouse model data was sourced from GSE102556[27]. Further, the expression changes seen in CVR are likewise reverse of those driven by Li. see Table 1. The combined BP profile shows a smaller but still significant anti-correlation with ALP, see Table 1. This opens up the interesting possibility of validating intervention candidates in the mouse by tracing the expression changes driven by therapeutic candidates.

**Figure 2.**
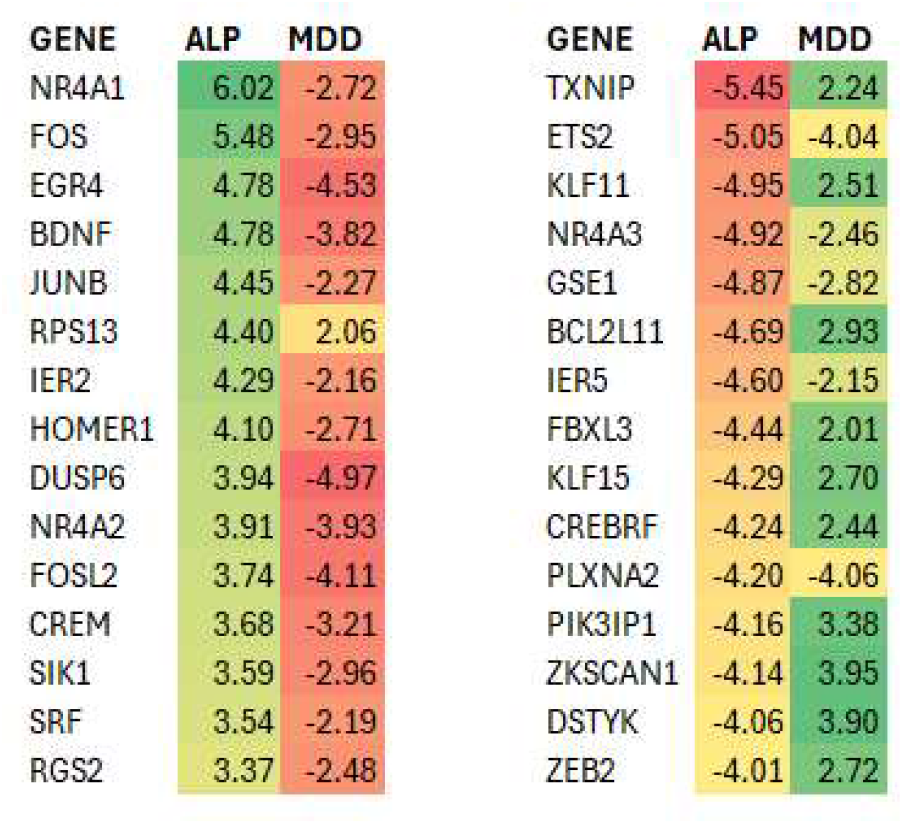
Li shows substantial reversal of expression changes in MDD and BPD. The topmost Li regulated genes are shown together with the expression changes in the MDD (left) and BPD (right) showing a clear reversal. Of the 40 most regulated genes due to Li treatment 32 in 40 and 22 in 32 are regulated in an opposite sense in MDD and BPD.

**Table 1.** There is a consistent anti-correlation of the acute Lithium response profile and gene expression changes seen in MDD BPD and a mouse model of depression. There is a significant anti-correlation of the acute Lithium profile with the combined MDD and BPD profiles, the correlations are given as Pearson correlation coefficients with the 95% confidence intervals and associated null probabilities. Interestingly a mouse model of depression shows similarities with both the MDD and BPD profiles as well as a significant anti-correlation with the Lithium profile.

| PROFILE | Lithium | MDD COMBO | BPD COMBO |
| --- | --- | --- | --- |
| MDD COMBO | -0.24 [-0.34 -0.13] 2.65e-05 |  |  |
| BPD COMBO | -0.13 [-0.23 -0.03] 0.01133 | 0.21 [0.12 0.29] 8.18e-06 |  |
| CUS Mouse | -0.25 [-0.32 -0.17] 2.46e-09 | 0.38 [0.30 0.46] <2.2e-16 | 0.13 [0.05 0.21] 0.001 |

### Repurposing candidates

Transcription provides a rational route to repurposing with the hypothesis that drugs reversing the gene expression changes associated with disease states may ameliorate the condition[13, 14, 18]. Further, provided that an existing drug’s activity is encoded in transcription[14, 23], this can facilitate a target for sourcing alternative therapeutics. As shown above, both Li treatment of cortical cultures and the MDD depression state in the brain are associated with substantial transcriptional changes. Further, bolstering the anti-depressive effects of Li, the expression changes driven by Li are significantly anti-correlated with those seen in MDD brain derived samples. This led us to adopt a transcription-based repurposing strategy for Li and look for alternative drugs with transcription profiles correlating with the Li profile and also showing an anti-correlation with the MDD expression changes. To this end we performed queried the CMAP set of 1,309 drug-like compound driven expression profiles. As can be seen in Table 2, we found eight cardiotonic steroids both significantly correlate with the Li profile and anti-correlate with the MDD summary profile.

**Table 2.** Cardiotonic steroid expression signatures correlate with the Li profile and anti-correlate with a summary MDD profile. The correlation Z scores for seven cardiotonic steroids from CMAP against the Li and MDD summary profile are shown together with the relevant rank (positive and negative respectively).

| COMPOUND | Lithium |  | MDD |  |
| --- | --- | --- | --- | --- |
|  | Z | RANK | Z | RANK |
| digitoxigenin | 5.22 | 12 | -5.73 | 14 |
| digoxin | 5.2 | 13 | -6 | 9 |
| ouabain | 4.27 | 21 | -7.19 | 2 |
| proscillaridin | 4.26 | 22 | -5.27 | 24 |
| digoxigenin | 3.82 | 35 | -2.99 | 168 |
| lanatoside C | 3.43 | 46 | -6.19 | 6 |
| helveticoside | 2.74 | 94 | -4.95 | 31 |

**Table 2.** CTS prescription is consistently associated with a lower Li prescription likelihood. The log odds and statistics are shown for the full NorPD data for those on a psychiatric medication over the period 2010 to 2021. CTS use negatively correlates with Li prescription in the full cohort (ALL), in the cohort with prescription status assigned to those with 10 or more prescriptions for the drugs, in age restricted cohorts and separately for men and women.

| Cohort | LogODDS [95% COI] | prob | Li and CTS | Li | CTS | TOT |
| --- | --- | --- | --- | --- | --- | --- |
| ALL | -1.15 [-0.94 -1.37] | 6.24E-27 | 88 | 17488 | 33234 | 1926252 |
| Pr > 10 | -0.96 [-0.63 -1.28] | 6.82E-09 | 37 | 10607 | 15784 | 1901972 |
| > 40 years | -0.96 [-0.75 -1.17] | 4.55E-19 | 88 | 14231 | 33177 | 1525892 |
| > 70 years | -1.03 [-0.78 -1.29] | 2.61E-15 | 60 | 4355 | 30290 | 652988 |
| men | -1.18 [-0.88 -1.47] | 1.07E-14 | 44 | 7663 | 15155 | 802285 |
| women | -1.14 [-0.84 -1.44] | 6.06E-14 | 44 | 9825 | 18079 | 1123967 |

### Prescription frequencies of CTS and Li in the NorPD psychiatric medication cohort

We have motivated the repurposing of CTS as alternatives to Li treatment in severe depression based on shared features in gene expression perturbation. It is then of interest to see to what extent prescription of CTS is associated with a reduced likelihood of being prescribed Li. To focus on the specific factors leading to Li prescription we decided to restrict our analysis to those being prescribed at least one form of psychiatric medication, see Methods for medications. This cohort comprises a significant fraction of the entire population (2 million of 5.5 million). To this end we compiled prescription records for 94 psychiatric medications and two CTS (digoxin and digitoxin) for the entire Norwegian population covering the years 2010 to 2021, see Methods. As can be seen from Table XXX there is a reduced Li prescription frequency in those prescribed cardiotonic steroids. Performing a logistic regression with age and sex as covariates we obtain a log odds association of Li with cardiotonic steroid use of -1.15 (95% CO[-0.94 -1.37] p = 6.24E-27). Here, medication status is binary with prescription status assigned to those with at least one prescription of the given medication. To test the robustness of the negative association we varied the minimal prescription count required for positive prescription status, see table XXX. We also varied the cohort minimal age and restricted analysis based on sex. We observed a consistent negative association of cardiotonic use with Li prescription, see table XXX. This observation bolsters our transcriptional analysis.

## Conclusions

The compromising side-effects and poorly understood mechanism of action of the widely prescribed Li in the context of major depression motivated our investigation into the possibility of repurposing drugs with established safety profiles that might be alternatives to Li treatment. We sought to delimit the Li activity mechanism through an analysis of acutely driven gene expression changes in the context of Li treatment of neuronal cells. Our lithium profile, ALP, showed an upregulation of key genes involved in neuronal function. Notably, brain derived neurotrophin, BDNF, genes involved in synaptic mitochondrial function, NR4A, and learning and memory, CAMKK2. We also found that ALP shows significant anti-correlation with human brain sample derived MDD and BP profiles. These observations led us to investigate the potential of ALP to capture repurposing candidates from databases of drug driven expression profiles. Querying CMAP with ALP we found that cardiotonic compounds constituted high rank hits. Interestingly cardiotonics also showed a significant anti-correlation with MDD and the mouse model CVS profiles. Bolstered by these observations we investigated whether prescribed CTS are associated with a lower Li prescription rate. Our analysis of the NorPD prescription data for those on psychiatric medication showed that CTS use is associated with a lower likelihood of Li prescription.

These observations motivate further investigation of the potential of CTS as an alternative to Li in the context of severe depression. However, the results presented here are only in the form of correlation analyses and do not establish a causal basis for the novel therapeutic potential of CTS. It would be of interest, for example, to see to what extent CTS reverses behavioural changes in the CVS mouse model of depression. The epidemiological analysis showing a decreased prescription of Li in those on CTS medication goes some way in validating CTS as a candidate. Such retrospective observations though have sometimes led to disputed claims, such as the proposed protective effects of statins in cancer[42]. A retrospective analysis of NorPD prescription data provided strong evidence for the protective effects of salbutamol against PD[24]. Subsequent analysis of United States Medicare data controlling for predicted smoking incidence found that the negative salbutamol association with PD incidence no longer obtains[43]. Their argument being that salbutamol use is higher in smokers and smoking reportedly lowers PD risk[44]. Interestingly, a transcriptional approach to repurpose drugs on the basis of their driving growth factor expression in the brain and subsequently providing trophic support for neurons in the context of PD led to the identification of salbutamol as a potential candidate, which was shown to reverse dopaminergic loss in a mouse model of PD[21].

It has been reported that heart disease can be a co-morbidity of clinical depression, with cardiovascular mortality being higher in those with depression[45]. This can be explained on the basis of lifestyle changes associated with depression or could have a more direct physiological connection as both conditions can be characterised by increased inflammation[46, 47] and autonomic nervous system dysregulation may lead on to cardiac problems[48]. These observations would point towards a higher prescription rate of heart medications in the major depression cohort and it would be of interest to contrast the prescription rates of CTS with other heart medications. In conclusion, our analysis suggests a possible role for CTS as a Li replacement in MDD and motivates further investigation in the form of a controlled trial.

## Acknowledgements

This work was supported by Medical Research Council grant MR/M013944/1, awarded to RK.

## Ethical approval

The project obtained approval from the regional ethics committee in Norway (REK ID 424276) and was evaluated to comply with the data protection regulations (SIKT ID 142811). Since the project uses data from national health registers, the project also has approval from the Directorate for e-health regarding exceptions from confidentiality (H-595 DOC-40609). All variables have been reviewed and assessed regarding data minimisation, relevance, and necessity in relation to achieving the research objective, cf. Section 32 of the Health Research Act.

